# A Musculoskeletal Model of the Hand and Wrist Capable of Simulating Functional Tasks

**DOI:** 10.1101/2021.12.28.474357

**Authors:** Daniel C. McFarland, Benjamin I. Binder-Markey, Jennifer A. Nichols, Sarah J. Wohlman, Marije de Bruin, Wendy M. Murray

## Abstract

**Objective:** The purpose of this work was to develop an open-source musculoskeletal model of the hand and wrist and to evaluate its performance during simulations of functional tasks.

**Methods:** The musculoskeletal model was developed by adapting and expanding upon existing musculoskeletal models. An optimal control theory framework that combines forward-dynamics simulations with a simulated-annealing optimization was used to simulate maximum grip and pinch force. Active and passive hand opening were simulated to evaluate coordinated kinematic hand movements.

**Results:** The model’s maximum grip force production matched experimental measures of grip force, force distribution amongst the digits, and displayed sensitivity to wrist flexion. Simulated lateral pinch strength fell within variability of *in vivo* palmar pinch strength data. Additionally, predicted activation for 7 of 8 muscles fell within variability of EMG data during palmar pinch. The active and passive hand opening simulations predicted reasonable activations and demonstrated passive motion mimicking tenodesis, respectively.

**Conclusion:** This work advances simulation capabilities of hand and wrist models and provides a foundation for future work to build upon.

**Significance:** This is the first open-source musculoskeletal model of the hand and wrist to be implemented during both functional kinetic and kinematic tasks. We provide a novel simulation framework to predict maximal grip and pinch force which can be used to evaluate how potential surgical and rehabilitation interventions influence these functional outcomes while requiring minimal experimental data.

## I. Introduction

COMPUTATIONAL musculoskeletal models of the hand and wrist provide valuable insight into how hand dysfunction occurs following changes to specific musculoskeletal structures [1], neural control signals [2], and functional use [3]. However, due to the complexity of the hand and limited physiological data characterizing the middle, ring, and little fingers, most hand models are used in simulations involving only the thumb and/or index finger [1, 2, 4-12]. As a result, more complex functional tasks that require coordinated effort from all the digits (e.g., grasping, hand opening) have rarely been simulated. Currently, simulation studies of hand function tend to focus on fingertip/pinch force [1, 2, 8-10] or kinematic motion involving a single digit [2, 5, 6].

Recently, there have been efforts to develop more complete musculoskeletal models of the hand and wrist [13-17]. These models advance the field in that they include all the digits and muscles of the hand. However, only one research group reports implementing their model in simulations of grasping [3, 14, 18], using inverse-dynamic methods. Specifically, experimental data quantifying joint posture, contact points, and contact forces for all the digits of the hand served as inputs to a musculoskeletal model that included the wrist, all digits, and all muscles to evaluate muscle and joint loading during grasping [3, 14, 18]. Despite the value of this work, inverse-dynamic methods are not an ideal method to study muscle coordination of movement and require assumptions regarding how net joint torques are produced by multiple muscle forces. In contrast, forward-dynamics represents the way the body processes neuromuscular excitation signals to produce movement [19]. In particular, optimal control theory, which uses a forward-dynamics simulation to optimize activations to accomplish a hypothesized task, has been posited to have more potential to provide insight than inverse-dynamics simulations when examining why a particular muscle coordination pattern is chosen to accomplish a task [19]. For example, optimal control theory has been shown to better replicate muscle coordination during cycling than static optimization [20]. Additionally, optimal control theory has the potential to accurately predict maximal performance of specific tasks with minimal experimental data. For example, optimal control theory has been used to simulate a maximum-height squat-jump, and this optimization reproduced the major features of experimental data including the ground reaction forces, order of muscle activity, and overall jump height [21].

Several modeling challenges exist when developing musculoskeletal models of the hand and wrist. These challenges span multiple areas, including representing the complex motion of the wrist bones [22, 23], simulating force transmission via the extensor mechanism [24], defining passive joint stiffness parameters which are essential for simulating kinematic movement [2, 25], and incorporating moment arms and force-generating parameters for muscles where there is limited experimental data describing their capacity. It will take years and the cumulative contributions of multiple research groups to fully address all these challenges. Thus, there is a need for an open-source model to serve as a foundation for future work and to promote collaboration. Importantly, due to the scarcity of simulations involving coordinated functional tasks, we currently do not fully understand which modeling challenges are primary barriers to the field. The objectives of this work are 1) to develop an open-source model of the hand that includes the wrist, all digits and muscles of the hand, and passive joint properties for each flexion/extension degree of freedom, 2) to demonstrate, evaluate, and share the implementation of our model for simulations of maximal grip force, maximal lateral pinch force, active hand opening, and passive grasp and release (tenodesis [26]), 3) to describe how current modeling limitations influence the ability to simulate coordinated functional tasks, and 4) to identify key next steps to address the primary barriers to robust simulations of coordinated functional tasks. Our musculoskeletal model and simulation tutorials are freely available for download on simtk.org to enable others to build upon this work.

## II. Model Development

A dynamic musculoskeletal model of the hand and wrist was developed in OpenSim (v4.3) [27] by adapting and expanding upon existing simulation work completed by our group [1, 2, 6, 10, 28-32]. The musculoskeletal model implemented here includes 22 rigid bodies, with mass and inertial properties for the individual bone segments as described previously [6]. Also as described previously [29, 30], the kinematic model from Holzbaur *et al*. [31] was augmented to include experimentally-derived kinematics of the middle, ring, and little finger [33]. The current model includes 23 independent degrees of freedom (DOFs) including a flexion/extension DOF for each interphalangeal (IP) joint of the four fingers and thumb, flexion/extension and ab-adduction DOFs for each metacarpophalangeal (MCP) joint of the fingers, a flexion/extension DOF for the MCP joint of the thumb, flexion/extension and ab-adduction DOFs for the carpometacarpal (CMC) thumb joint, a coupled flexion DOF for the CMC joints of the ring and little finger, and flexion/extension and radial/ulnar deviation DOFs for the wrist.

The model includes passive joint properties for all flexion/extension DOFs of the phalanges and thumb, for CMC ab-adduction of the thumb, and for wrist flexion and deviation DOFs. Passive joint properties for the fingers and thumb DOFs were implemented as position-dependent torques [2, 5, 6]. Passive properties for the thumb and index finger were implemented from the literature [25, 34-36], as described in previous work [1, 6]. For the MCP joints of the middle, ring, and little finger, these torques were defined to match newly available experimental data [37] and added to the model. Data were not available for the proximal interphalangeal (PIP) and distal interphalangeal (DIP) joints of the middle, ring, and little fingers. Thus, the position-dependent torques were implemented as scaled versions of the position-dependent torques for the index finger. As described in Saul *et al*. [32], passive joint properties for the wrist were implemented as coordinate limit forces [38].

Forty-three Hill-type muscle-tendon actuators representing the intrinsic muscles of the hand, the extrinsic muscles of the hand, and the primary wrist muscles were included in the model. Muscle-tendon paths for the intrinsic muscles of the phalanges were added to the model to match experimental moment arms of MCP flexion [39]. Because MCP abduction moment arm data for the middle, ring, and little fingers does not currently exist, MCP abduction moment arms for these digits were modeled to be similar to MCP abduction moment arms for the index finger [40]. Muscle-tendon paths for the intrinsic thumb muscles, the extrinsic index finger muscles, and the primary wrist muscles, were implemented as specified in previous models [6, 10, 32]. Muscle-tendon paths for the extrinsic muscles of the middle, ring, and little fingers were implemented from Saul *et al*. [32], but were edited to match experimental moment arm data for MCP, PIP, and DIP joints [17, 39-41] since the original definition of these muscle paths did not include these DOFs [32]. The extensor mechanism was not modeled here; the intrinsic muscles inserted onto the proximal phalange, crossing only the MCP joint [13, 42]. The extrinsic muscles inserted onto the distal phalanges, crossing both interphalangeal joints.

As described in Binder-Markey and Murray [6], the “Millard2012EquilibriumMuscle” muscle model [43] with the active force-length, force-velocity, passive-force length, and tendon force-strain curves adjusted to replicate the respective curves in the Saul *et al*. model [32] were used for each muscle-tendon actuator. Muscle force-generating parameters including physiological cross-sectional areas (PCSA), optimal fiber lengths, and pennation angles were added to the model for the intrinsic muscles of the fingers [44, 45]; parameters defined in previous models were replicated for the remaining muscles [6, 10, 32]. Peak isometric forces for the intrinsic finger muscles were calculated from the PCSA values using a specific tension of 50.8 N/cm^2^, consistent with previous models [6, 10, 32]. Peak isometric forces for the primary wrist and extrinsic finger muscles were based upon *in vivo* muscle volume [46] and isometric strength [47] data of healthy young adult males. Thus, we intend the model to represent strength of healthy young adult males in the subsequent simulations.

Tendon slack lengths for the intrinsic muscles of the phalanges and the extrinsic finger muscles of middle, ring, and little fingers were calculated from muscle-tendon lengths and fiber lengths using the following equation: 

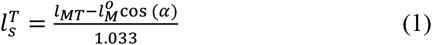

where *l*_*MT*_ is muscle-tendon length when all joints are in neutral position and the muscle is inactive, 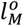 is the optimal fiber length, and *α*is pennation angle [2, 48]. Tendon compliance for the lumbricals was neglected (i.e. *l*_*ts*_=0) to improve simulation stability since this muscle-tendon unit had a small ratio of tendon slack length to optimal muscle fiber length [43]. Tendon slack lengths for the primary wrist muscles, extrinsic and intrinsic thumb muscles, and the extrinsic index finger muscles were implemented as specified in previous models [6, 10, 32].

## III. Kinetic Simulations

### A. Grip Strength

Grip force was computed using an elastic foundation contact model [49, 50] between the skin of the phalanges (massless cylinders overlaid on the bone geometries) and an elliptical cylinder representing a widely used dynamometer (Fig.1.A). Because the American Society of Hand Therapists recommends that the weight of the dynamometer be lightly supported during clinical grip strength measurements [51], we defined the cylinder representing the dynamometer to be massless as well. The diameter of the cylinder (48mm) was defined to represent setting II, the dynamometer setting typically associated with maximum strength [52]. The orientation and location of the dynamometer in the hand was confirmed in one subject using a handheld goniometer (Fig.1.B). For most adults holding the dynamometer on setting II, only the proximal and intermediate phalanges create contact force against the instrumented portion of the dynamometer (Fig.1.B), where the force component normal to the instrumented surface is measured as grip force. To replicate this instrumentation, only contact forces from the proximal and intermediate phalanges normal to the major axis of the elliptical cylinder contributed to simulated grip force. Whereas contact between the distal phalanges and the elliptical cylinder did not contribute to simulated grip force, contact surfaces on the distal phalanges were included so they did not cross through the surface of the elliptical cylinder during the simulation. The elliptical cylinder was attached to the distal end of the third metacarpal with a weld joint, preventing slip. Because the dynamometer is only instrumented at the fingers, the thumb was locked during all simulations and the 5 intrinsic thumb muscles were excluded from all grip force simulations, as they would not contribute to either joint motion or force production. Contact parameters representing the skin and the dynamometer were taken from the literature [53-55] (Table I).

**Table I:** Contact Parameters

|  | Young's modulus | Poisson's Ratio | Coefficient of friction |
| --- | --- | --- | --- |
| Skin | 12.3 Mpa | 0.48 | 0.29 |
| Dynamometer | 1.75 GPa | 0.43 |  |

**Fig. 1.**
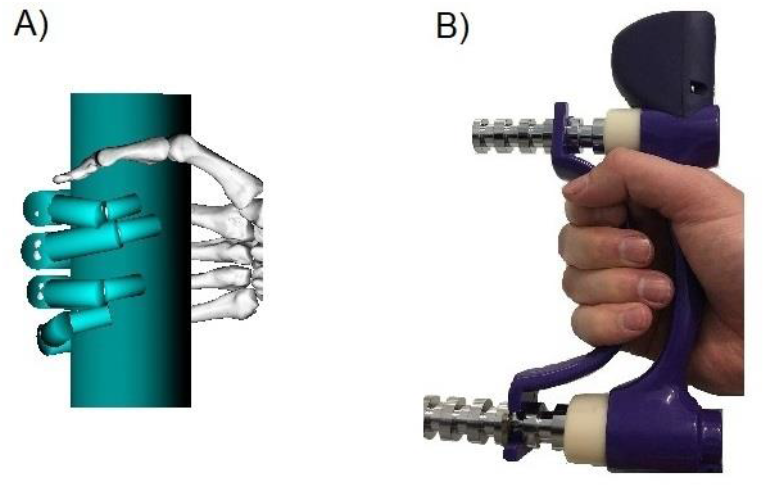
A) Our model’s representation of grasping a standard dynamometer on setting II. Contact bodies (teal cylinders) are overlaid on bone geometries. Only contact from the proximal and intermediate phalanges contribute to grip force to mimic a person holding the dynamometer (B).

Optimal control theory was implemented to simulate grasping the dynamometer. A set of 15 independent muscle activations were optimized to maximize the contact force normal to the major axis of elliptical cylinder while maintaining an initial wrist posture (Top of Fig. 4). The simulated-annealing optimization (MATLAB R2018b, The MathWorks Inc., USA) would alter the set of input activations and run the forward-dynamic simulation via the OpenSim API for each step of the optimization. The timescale of each simulation was 0.15s. Each individual muscle activation was constrained to a constant value throughout a given simulation; preliminary simulations did not show improved performance when using discrete activation nodes. During the forward-dynamic simulation the wrist was not constrained, although the optimization encouraged the wrist to maintain the initial posture with a function that penalized wrist movement. Maximal grip force was defined as the average contact force during the simulation. Grip force generally reached a constant value within 0.01 seconds, the simulation time was extended to ensure the model maintained the initial wrist posture.

For 12 muscles (the primary wrist muscles, the extrinsic thumb muscles, extensor indicis proprius and extensor digiti minimi), independent activation levels were defined for each muscle. Three additional independent activation levels were defined, one each for the flexor digitorum profundus (FDP), flexor digitorum superficialis (FDS), and extensor digitorum communis (EDC). These 3 multi-compartment muscles are each represented in the model with 4 muscle “slips” that actuate each of the fingers, comprising 12 muscle-tendon actuators in the model. We chose to define a single activation level for each set of 4 muscle “slips” because experimental work shows that voluntary activation of the middle compartment of FDP and EDC cause similar involuntary co-activation in the index compartment of the same muscle during isometric force production [56].

Muscle activations for the remaining 14 muscle-tendon actuators (43 total less 12 single compartment, 3×4 multi-compartment muscles, and 5 intrinsic thumb muscles) were manually defined to limit the solution space. Specifically, the 14 intrinsic muscles for the index, middle, ring, and little fingers were set to full activation. Because the extensor mechanism was not modeled, these intrinsic finger muscles create flexion moments about the MCP joints without extending the distal phalanges. Therefore, activation of these muscles always both increased contact force and contributed to the simulated grip force.

The optimization to determine simulated grip force was run 50 times since simulated annealing does not guarantee that the global optimum will be determined in a given iteration. We report the average and standard deviation for grip force, distribution of force amongst the digits, and predicted activations for the 5 simulations with the best objective function values. For these simulations, the wrist was set in extension with slight ulnar deviation (Fig. 4: Self-selected posture) to replicate the experimental wrist posture associated with maximum grip strength [57]. Initial postures of the joints in the hand were selected such that the contact cylinders representing the skin and the dynamometer were in contact at the start of the simulations. While not included as DOFs in the model, the hand and wrist model was connected to a previously described model of the upper limb [32], the shoulder was positioned in neutral abduction, the elbow was in 90° flexion, and the forearm was in neutral pronation/supination; this is recommended arm posture for clinical grip strength measurements [51]. Simulation results were compared to experimental studies in which grip strength was measured [57-59], as well as descriptions of normative grip strength data of nonimpaired young adult males [60-63]. Distributions of force among the individual digits were evaluated and compared with experimental distributions of finger force during grip [58]. Additionally, optimized activations were compared to electromyography (EMG) data reported in the literature [59, 64]. Lastly, final wrist postures of the simulations were evaluated to confirm that the wrist posture did not drastically change during the simulations. We repeated maximum grip force simulations in multiple wrist postures (see top Fig. 4 for postures) to compare with experimental data that describes the influence of posture on grip strength [57].

### B. Pinch Force

The same optimal control simulation framework was used to determine a set of muscle activations that maximizes lateral pinch strength. As we have implemented previously [1, 65], pinch force was defined as average constraint force in the palmar direction of the global coordinate frame between the model’s ground frame and a massless body welded to the thumb-tip. Using a penalty term, off-axis forces in the medial-lateral and proximal-distal directions were required to be less than 17% of the palmar force to mimic the experimental methods of Valero-Cuevas *et al*. [9].

The model was set in a lateral pinch posture (15° CMC extension, 20° CMC adduction, 20° MP flexion, and 40° IP flexion) with 0° wrist flexion and 0° wrist deviation [1, 65]. The forward-dynamics simulation methods mean the model can move from the initial posture to the equilibrium posture that results from muscle force production about multiple joints. Due to the constraint between the ground frame and the thumb-tip, this final thumb posture always remained consistent with a lateral pinch. During the optimization the wrist was not constrained, although a penalty term encouraged maintenance of the initial posture within 5° in any direction. The DOFs for the index, middle, ring, and little fingers were locked during the simulation.

The optimization solved for 15 independent muscle activations of the intrinsic thumb muscles, the extrinsic thumb muscles, and the primary wrist muscles. The intrinsic finger muscles (14 muscle tendon-actuators) and the extrinsic finger muscles (14 muscle tendon-actuators) were removed from these simulations.

## IV. Kinematic Simulations

### A. Hand Opening

Static optimization [66] was implemented to predict muscle activations for the sign language letter “O”; this motion was chosen since the motions involved are similar to hand opening and closing (Fig 2). Kinematics were collected on two subjects with a Cyberglove II motion capture glove (Cyberglove system LLC; San Jose, CA). This motion capture system is a “one-size fits-all” glove with 22 resistive bend sensors that record at 90Hz. The raw data is converted into joint angles by applying sensor gains determined during specific calibration tasks [30]. Joint angles were filtered with a 3^rd^ order Butterworth low-pass filter with a cutoff frequency of 6Hz, and the average kinematics of the two subjects were used in the simulation. Reserve torque actuators (max torques 0.1Nm) were included for the PIP and DIP DOF for the index, middle, ring, and little fingers, CMC flexion and abduction of the thumb, and coupled flexion of the CMC joints of the ring and little finger.

**Fig. 2.**
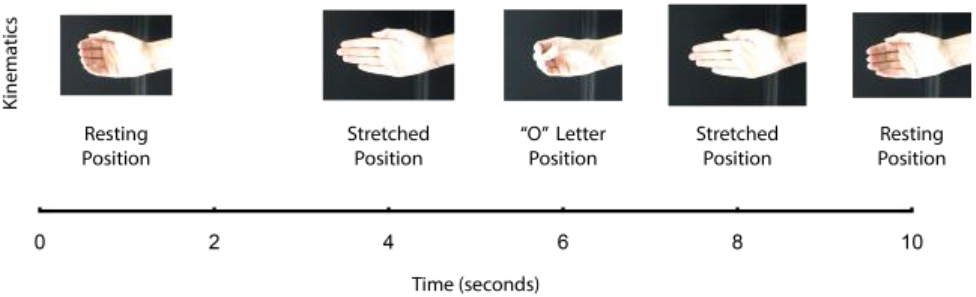
Kinematic motion of sign language letter “O”.

Predicted activations from the static optimization were compared with the average electromyography (EMG) signals of our two subjects. During the kinematic data collection, EMG data of extensor pollicis longus (EPL), extensor pollicis brevis (EPB), abductor pollicis longus (APL), flexor pollicis longus (FPL), flexor pollicis brevis (FPB), adductor pollicis (ADP), EDC, and FDS were collected with fine-wire electrodes with Delsys Bagnoli-16 system (Delsys Incorporated, Natick, MA) at 2000Hz. Electrode insertion points were identified using an ultrasound system (Siemens Medical System Inc., Malvern, PA) with a 4.5 cm linear array probe. Using a 27-gauge hypodermic needle, bipolar fine-wire electrodes were inserted in each muscle and electrode placement was verified by checking muscle activity data during standard manual muscle testing postures. Subjects performed a series of isometric maximal voluntary contractions to normalize EMG measurements [48]. Raw EMG data was post-processed by band-pass filtered (25-500Hz), notch filtered (59.5-60Hz) to remove power line noise, rectified, and low-pass filtered at 8Hz (4^th^ order recursive Butterworth filters). Data was then Gaussian smoothed with a 100ms window and normalized to their respective MVC peak. The human subjects protocol was approved by the Institutional Review Board (IRB) of Northwestern University (IRB Study: STU00039072; initial approval 1/7/2011); participants gave informed consent prior to participation.

### B. Tenodesis (Passive Grasp and Release)

Passive simulations of tenodesis grasp and release were performed [6]. In this forward-dynamics simulation, all muscle-tendon actuators were included and held at 0 activation for the duration of the simulation while wrist motion was prescribed. To simulate tenodesis grasp, wrist posture was maintained at a posture of 60° flexion for 1s to yield an initial equilibrium posture for the digits and then wrist extension was prescribed at 20°/s until the wrist achieved 60° extension. To simulate tenodesis release, wrist posture was held at 60° extension for 1s and then wrist flexion was prescribed at 20°/s until the wrist achieved 60° flexion. During the simulation, all flexion/extension DOFs for the index, middle, ring, and little fingers were unconstrained and simulated with time; the remaining finger and thumb DOF were locked. Kinematic motion was evaluated to confirm that the model exhibited coupled movements between the wrist and phalanges that resulted in a grasping posture during wrist extension and an extended posture during wrist flexion.

## V. Results

### A. Grip Strength

Simulated maximum grip force was consistent with reported grip strength from several experimental studies [57-59] (Fig 3.A) and fell within the range of reported normative grip force (36.2-54.9kg) for healthy adult males between ages of 20 and 30 [60-63]. Notably, both wrist posture (Fig 3.D) and participant demographics are inconsistent between available experimental studies that report grip strength. Kong *et al*. [58] and Mogk and Keir [59] included young adults (males only in [58]; males and females in [59]) whereas O’Driscoll *et al*. [57] include young and middle-aged males and females. Using the ‘self-selected’ wrist posture identified in O’Driscoll *et al*. [57], the model’s grip force was 44.5±0.7kg; experimental grip strength in this posture was 41±13.4kg.

**Fig. 3.**
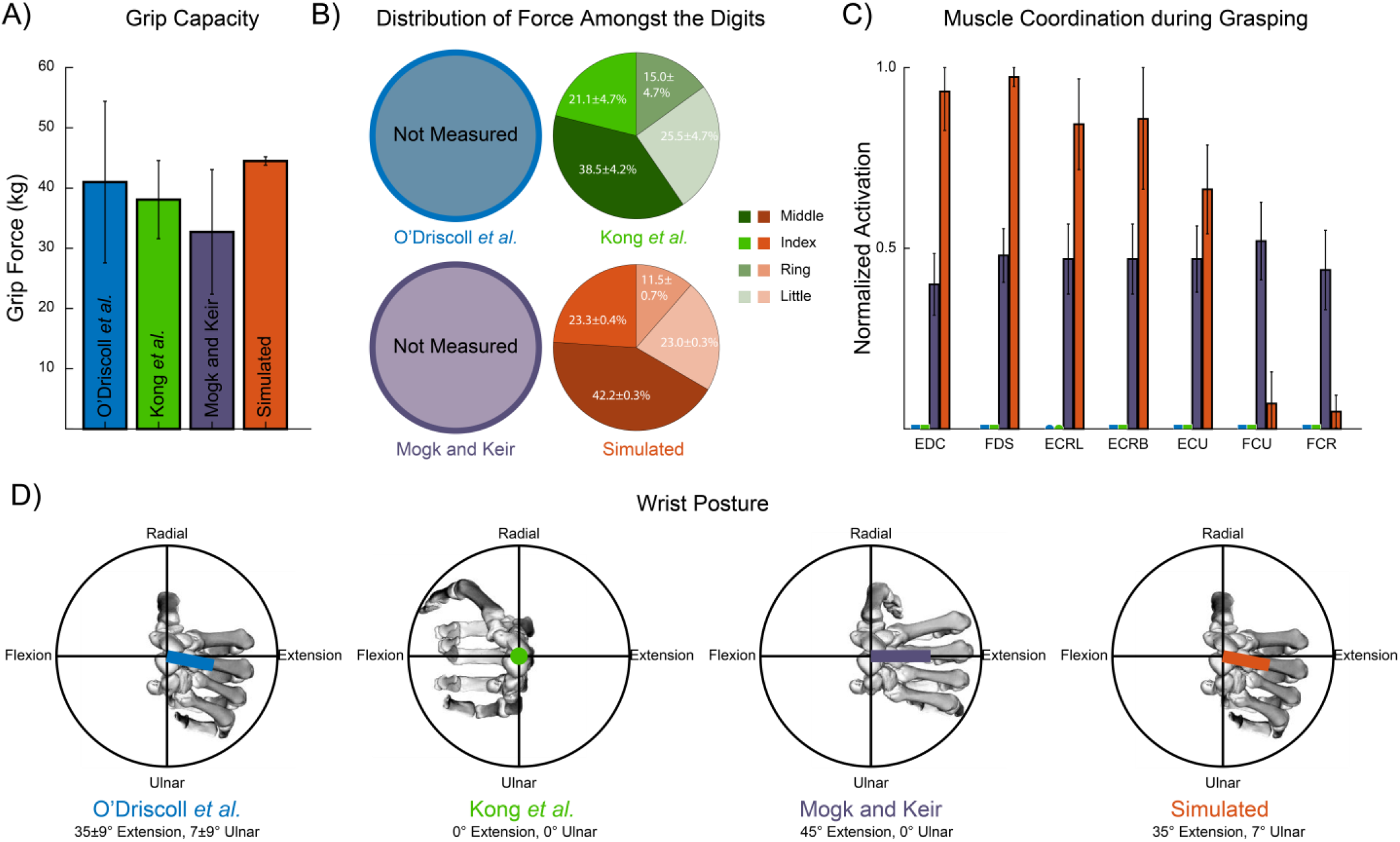
A) Average and standard deviation of experimentally measured maximum grip force, as reported from several studies (blue, green, and purple bars) and the grip force from the 5 simulations with the best objective function values in ‘self-selected’ wrist posture (orange bar). B) Average and standard deviation of the distribution of force amongst the individual digits from the same studies and simulations; of these studies, only Kong *et al*. (2011) reported the distribution of force amongst the digits. C) Average and standard deviation of muscle activations from the same studies and simulations; of these studies, only Mogk and Keir (2003) report EMG signals measured during grasping (color coded bars for the other two studies are set to zero on this graph for consistency across panels). D) Illustrations of the wrist posture adopted during grip force, as reported by each experimental study and the wrist posture used for our simulation results. The overlaid color-coded vectors represent the magnitude and orientation of the initial wrist posture in a Cartesian coordinate system with the positive x-axis representing wrist extension and the positive y-axis representing radial deviation. The origin of this coordinate system is aligned with the base of the lunate.

The simulated grip force was comprised of a similar distribution of force production amongst the digits when compared to Kong *et al*. [58] (Fig 3.B). In the simulations, the middle finger contributed the most to grip force (42.2±0.3%), followed by the index and ring finger (23.3±0.4% and 23.0±0.3% respectively). The little finger produced the least force 11.5±0.7%.

The optimization predicted higher activations for extrinsic finger and wrist extensor muscles and lower activations for wrist flexor muscles than EMG data reported during maximal grip strength (Fig 3.C) [59]. The largest difference between EMG data and predicted activations was with the wrist flexors. In particular, flexor carpi ulnaris (FCU) and flexor carpi radialis (FCR) had low activations for the simulations (0.07±0.08 and 0.05±0.05 respectively). For the remaining muscles, (the three wrist extensors, EDC, and FDS), both the experimental study and our simulations indicate that intermuscular co-activation levels were relatively consistent despite the overall difference in magnitude of activation.

Among the 5 simulations with the best objective function values from our 50 repeated optimizations, there was generally larger variation in the simulated muscle activations than the simulated grip force. The coefficient of variation (CoV) for simulated grip strength was 0.02. Apart from FDS (CoV = 0.03), CoVs for muscle activations were at least an order of magnitude greater than grip force. Specifically, for the extensor (wrist and extrinsic finger) muscles’ CoV ranged from 0.12 to 0.23; CoVs for FCR and FCU was 0.95 and 1.25 respectively.

Whereas the model did exhibit a dependence of maximum grip strength on wrist posture, the simulated maximum grip force of the model did not replicate the specific sensitivity to wrist posture reported in O’Driscoll *et al*. [57] (Fig. 4). In O’Driscoll *et al*. [57], grip strength was always weaker when wrist posture was shifted away from the ‘self-selected’ posture in any direction (p<0.0001: paired t tests reported in [57]). In contrast, the model was strongest (45.4±0.6kg) when the wrist posture was shifted from the ‘self-selected’ posture in extension; the simulated increase in grip force was small but significant (p<0.05: one-way repeated-measures ANOVA). Among our simulations that replicated the wrist postures from O’Driscoll *et al*. [57], only the wrist posture shifted in flexion was significantly weaker than ‘self-selected’ posture in our simulations (p<0.05: one-way repeated-measures ANOVA); the decline in simulated grip force (4.9kg, ∼11%) was less substantial than observed experimentally (11kg; ∼27%) in this posture (Fig 4). Both of the 2 additional postures we simulated (neutral and flexed) were significantly weaker than the simulations performed in the experimental postures from [57] (p<0.05: one-way repeated-measures ANOVA); the declines in simulated grip force when compared to the ‘self-selected’ posture were 18.3kg (∼41%) and 20.4kg (∼46%) for the neutral and flexed wrist postures, respectively. All simulations maintained the initial wrist posture within 5° for each direction.

**Fig. 4.**
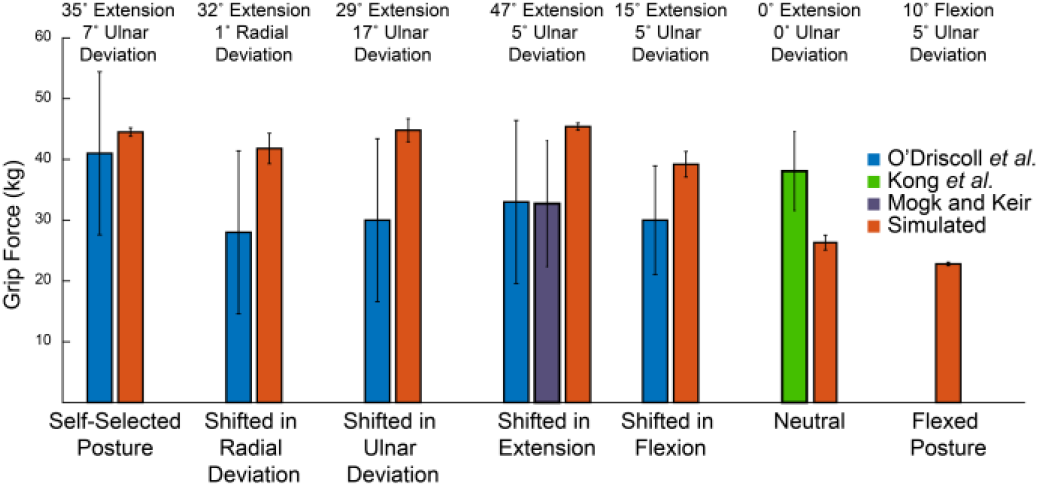
Simulated maximum grip force was compared to experimental measures in several wrist postures. Blue, green, and purple bars represent the experimental grip force reported in [57], [58], and [59] respectively. Error bars represent the standard deviation of the experimental data. Orange bars represented the simulated maximum grip force; average and standard deviation of the 5 simulations with the best objective function values (top 10%) are displayed.

### B. Pinch Force

Simulated lateral pinch force fell within variability of *in vivo* palmar pinch strength. *In vivo* palmar pinch force measured under similar conditions that limit off-axis forces is 51.9±20.4N [9]. Simulated pinch force of the top five simulations was 66.3±2.3N; all of these simulations maintained maximum off-axis force within 17% of the palmar force. Activations that maximized pinch force in our simulations for all muscles except abductor pollicis longus (APL) fell within the variability of normalized EMG data during palmar pinch force production [9] (Fig. 5). Lastly, the simulations maintained the initial wrist posture within 5°.

**Fig. 5.**
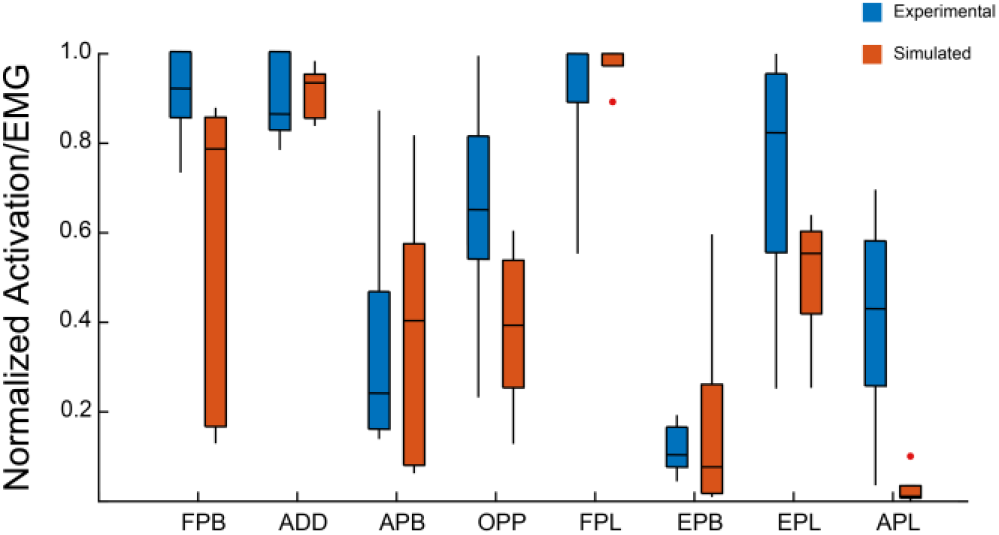
Simulated activations for the 5 pinch simulations with the best objective function values (top 10%) (orange boxplot) were compared to the range of normalized EMG data during palmar pinch (blue boxplot) [9].

### C. Hand Opening

Overall, predicted activations fell within 2 standard deviations of the normalized EMG signals for the hand opening task (Fig. 6). Additionally, timing of activation peaks generally aligned with the peaks in the EMG data (Fig. 6). Static optimization overpredicted activations for extensor pollicis brevis (EPB) on the second peak. Whereas predicted activations for flexor pollicis brevis (FPB) generally fell within 2 standard deviations of the normalized EMG data, the static optimization did not predict the large peak in activation seen in the EMG data. Instead, during this peak, the static optimization fully activated opponens pollicis (OPP). Predicted activations of EDC did not align well with the EMG data, particularly at the start of the motion.

**Fig. 6.**
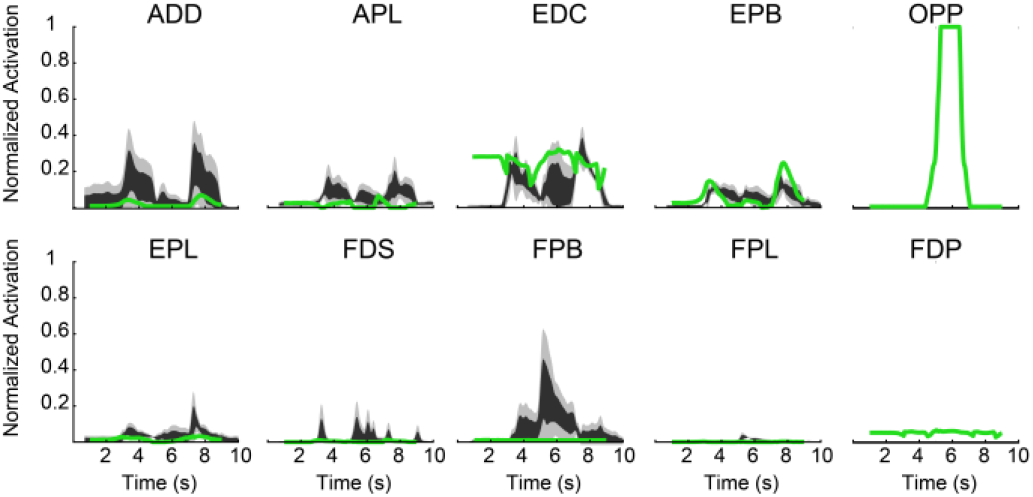
Simulated activations (green line) for the hand opening task generally fell within 2 standard deviations of experimental data (1 standard deviation: black region, 2 standard deviations: light grey region). OPP and FDP did not have EMG data available for comparison.

### D. Tenodesis (Passive Grasp and Release)

In the passive grasp and release simulation, the model displayed coupled motion between the wrist and fingers mimicking tenodesis. During prescribed wrist extension, the model passively flexed the digits creating a loose grasping posture (Fig. 7). Likewise, during prescribed flexion, the digits passively extended. On average, the MCP range of motion was 65.2°. On average, the PIP range of motion was 16.1°. The DIP joint flexion was constant throughout the motion. All digits displayed a similar range of motion, although the digits moved through this range of motion with different trajectories. The index finger was the least similar to the other digits with maximum differences in joint angle at a given instance of 49.2° and 14.6° for the MCP and PIP joints respectively.

**Fig. 7.**
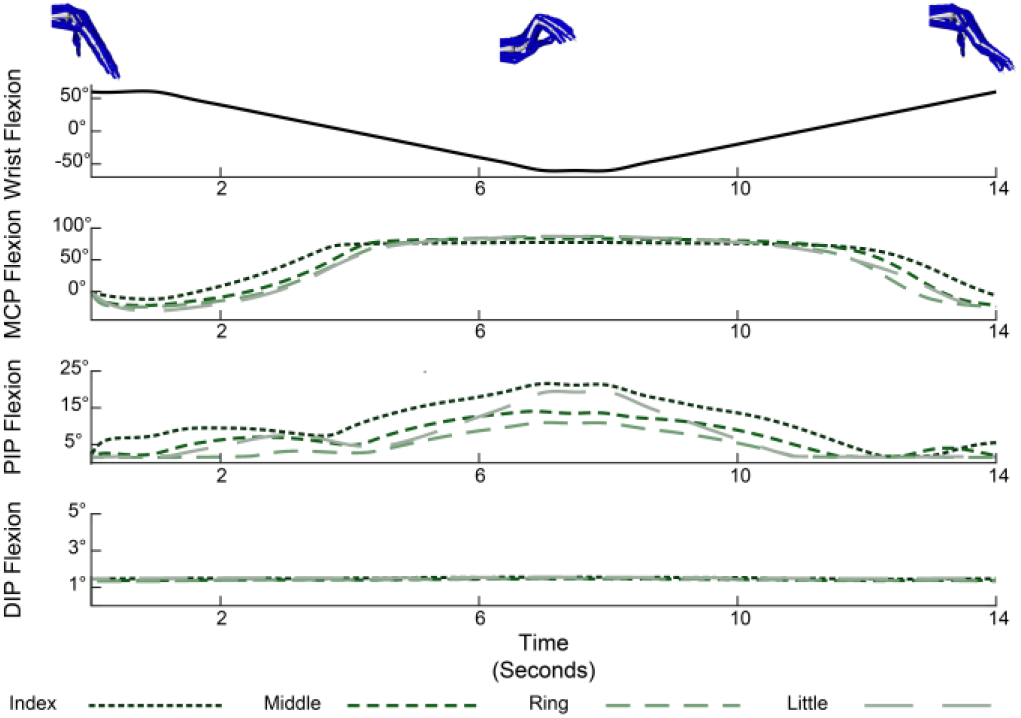
The passive simulation displayed the coupled motion between the wrist and fingers mimicking tenodesis grasp and release. The top panel displays the prescribed wrist flexion/extension. In order from top to bottom, the remaining panels display MCP, PIP, and DIP flexion/extension of the individual digits. Note differences in y-axis ranges.

## VI. Discussion

We have developed an open-source musculoskeletal model that includes all parameters necessary to perform muscle-driven, forward-dynamics simulations of force generation and multi-joint motion. Our model includes the wrist and all five digits of the hand, and is an extension (and compatible with) a previous open-source model of the shoulder, elbow, and wrist [32]. We provide examples of implementation of our musculoskeletal model for the simulation of maximal grip strength, maximal lateral pinch strength, passive hand opening and closing (i.e., the tenodesis grasp), and active hand opening. We also compare simulation results to experimental measurements to provide an assessment of the model’s performance. To simulate maximum grip and pinch strength, we developed an optimal control theory simulation framework that combines forward-dynamics simulations with a simulated-annealing optimization. This framework requires minimal experimental data to simulate maximum strength and is similar to common experimental maximum isometric strength protocols where participants are placed in an initial posture and instructed to produce the largest possible force. The kinematic simulations of passive tenodesis and active hand opening are also highly novel. Few prior studies have simulated coordinated hand motion with a model that includes the wrist and all five digits of the hand, and those that have do not evaluate functional tasks [67].

To our knowledge, no previously described musculoskeletal model of the hand and wrist has been implemented for muscle-driven, forward-dynamics simulations of both coordinated kinematic and kinetic functional tasks. Recently, several musculoskeletal models that include all the digits and major muscles of the hand have been developed [13-17]. However, few of these models are open-source and most are either not intended for or have not yet been shown to be usable for the full range of biomechanical tasks we describe here. For example, two models are not dynamic models; they do not include force-generating parameters for the muscle-tendon actuators and do not report the mass and inertial parameters for the rigid body segments [15, 16]. Whereas the remaining models are intended for dynamic simulations, implementation to simulate either grasp or pinch forces has not yet been described for two of the models [13, 17] (see [13, 17, 67] for more details on how these models have been used). The final model has been used in several inverse-dynamics simulations of grasping [3, 14, 18].

### A. Implementation of an optimal control theory framework for dynamic simulations of maximum grip and pinch strength

Currently, the few simulation studies of grip force that exist do so in an inverse framework [3, 14, 18]. Whereas inverse methods are useful for solving for muscle coordination patterns that can produce a specific force and are an important tool in the study of muscle coordination and joint loading, they have important limitations (e.g., identifying the cost function to solve the muscle redundancy problem, prescribing how forces are applied to each digit). Using optimal control theory, the model becomes representative of a research participant attempting to best complete a task and does not require assumptions about how the model will handle muscle redundancy or force production by the digits. We expect these methods will enhance the ability to perform ‘what-if’ simulations to evaluate how injury, disease, or surgical and rehabilitation interventions influence force production by the hand.

Whereas simulations of pinch and individual fingertip force production are more common, most of these simulations implement either an inverse or static framework [1, 8-10, 17, 42]. For example, several studies have used static optimization to determine muscle forces during fingertip force production [8, 17], but given the inverse framework, these simulations cannot readily predict maximum strength. For example, our lab group [1, 65] previously estimated maximum lateral pinch force using Computed Muscle Control simulations [68]. Whereas not entirely an inverse method (see [68] for details), Computed Muscle Control required the desired posture and pinch forces to be explicitly specified. Thus, to estimate maximum pinch force, we prescribed force in increasing 10N intervals, and interpreted a threshold force, beyond which the algorithm failed to identify muscle coordination patterns that produced greater forces, as an indicator of maximum strength. To predict maximal fingertip or pinch strength, other studies have used forward simulations; however, these simulations occurred in a mechanically static framework [9, 10, 42, 69]. For example, we previously replicated the methods of [9], to simulate lateral pinch with the same thumb model used here in a different computational environment [10, 69]. In this previous work, forward solutions were computed for the model in a prescribed, specific, static joint posture. In the simulations we present here, we define an initial posture, the optimization solves for a set of muscle activations, off-axis compensating endpoint forces, and resulting equilibrium posture that maximizes strength.

### B. Performance of model for force production

To evaluate our model, we compared the optimal strength simulations to published literature. Because there is no single data set available that describes maximum grip force [57-59], distribution of force amongst the digits [58], and EMG activity [59, 64], we made comparisons with multiple studies. The lack of a consistent data set measured in the same participants highlights an important gap in the field. For example, among the studies we compared to our simulations, wrist posture during testing and participant demographics varied considerably, and how these factors influence overall grip strength or the muscle coordination patterns used to generate force is not fully understood. In addition, the definition of the wrist posture used when quantifying grip force is not precise. From the anatomical definition of wrist range of motion, neutral wrist posture is defined as 0° of extension and 0° of deviation (e.g., [22, 70]). However, when grip strength is measured, clinical protocols specify a neutral wrist, in which the posture can involve wrist extension [71]. Thus, it is unclear the exact wrist posture used when wrist posture during grip measurements is reported as neutral without also specifying joint angles. Our model uses the anatomical definition of neutral wrist posture; we simulated grip strength in multiple postures to both address the ambiguity in the literature and the sensitivity of grip force to wrist posture.

Overall, our simulations compared well to grip force measured experimentally, with the distribution of simulated forces amongst the digits also consistent with experimental data [58]. Both the experimental data [59] and our simulations indicate that intermuscular co-activation levels amongst the three wrist extensors (ECRL, ECRB, and ECU), EDC, and FDS are relatively consistent. However, the optimization predicted higher activations for extrinsic finger and wrist extensor muscles and lower activations for wrist flexor muscles (Fig. 3). Despite these differences, overall, the optimal control theory simulations replicated many key features of maximal grip strength reported in the literature and provides a novel framework that can be combined with experimental work to better understand muscle coordination during grasping. For example, the difference in activation levels amongst primary wrist muscles between our simulation results and values reported in the literature suggest that additional functional criteria (e.g., stabilizing the wrist) are critical for grip force production. We anticipate both the model and the simulation methods we have developed here will play a role in future studies designed to answer the complex questions associated with understanding muscle coordination at the wrist and hand during force production.

The exact posture that maximizes grip force is debated [57, 72-74], but in general, experimental studies tend to agree that maximum grip force occurs with the wrist extended and with ulnar deviation. Our model was strongest in an extended and ulnar wrist posture, with grip force declining in more flexed postures. These results agree with data from Caumes *et al*. [71] that demonstrated modest declines in grip strength (<20%) as participants moved from their self-selected posture while still in wrist extension and larger declines (∼40%) in wrist flexion. Our strongest posture was more extended than the ‘self-selected’ posture described by O’Driscoll *et al*. [57] and our model did not display the same sensitivity to wrist posture reported in that study (Fig. 4). Because tendon slack length alters the relationship between joint angle and fiber length thereby influencing joint strength over the range of motion [6, 31, 75, 76], we analyzed the sensitivity of this result to our modeling choices for these parameters. Specifically, we re-defined the tendon slack lengths to be at their optimal length in the ‘self-selected’ grip posture (i.e. *l*_*MT*_ from equation 1 was determined in the ‘self-selected’ posture rather than with neutral wrist and fingers, see methods) and used the muscle activation patterns from the original optimizations with the adjusted model. While the model with the adjusted tendon slack lengths was stronger in each of the postures from [57], the sensitivity to wrist posture did not change (Fig. 8). While we did not re-optimize the muscle coordination strategies for all of the wrist postures, neither simulated grip force (cf. Fig. 8, open triangle) nor coordination patterns showed sensitivity to re-optimization with the adjusted model in the ‘self-selected’ posture. One interpretation of our sensitivity results is that sensitivity of grip strength to wrist posture may not be entirely due to biomechanical changes to force-generating capacity associated with wrist posture but may be due to changes in coordination to stabilize the wrist in un-ergonomic postures.

**Fig. 8.**
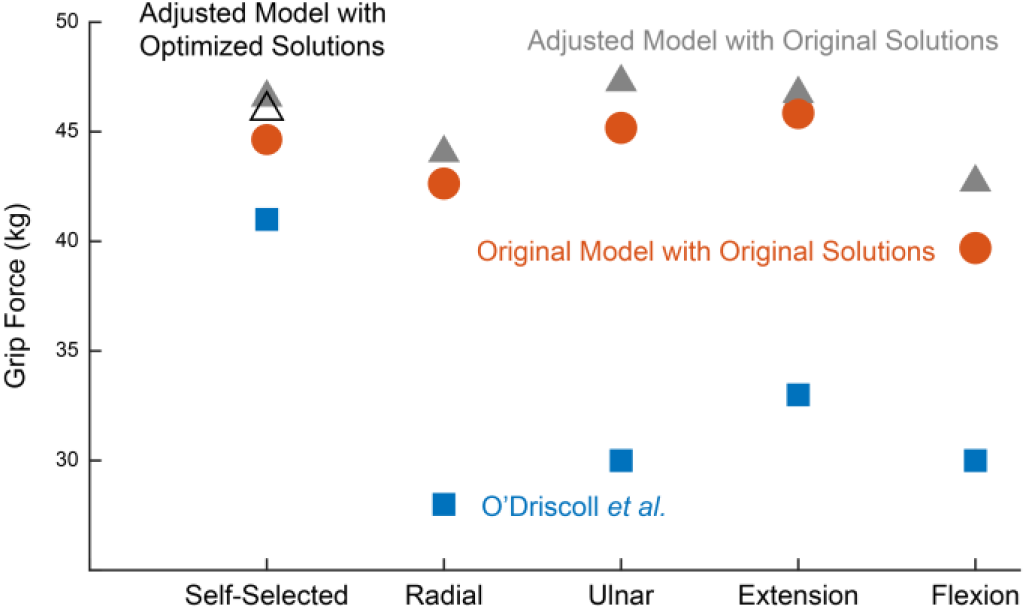
Sensitivity of grip strength simulations to tendon slack length. Adjusting the tendon slack lengths to be at optimal length in the ‘self-selected’ posture did not increase the sensitivity of simulate grip strength to wrist posture. Blue squares: experimental results from [57]. Orange circles: original simulation results from Fig. 4. Grey triangles: adjusted model with original muscle activation patterns. Black triangle: adjusted model with re-optimized muscle activation patterns.

We also compared the lateral pinch simulations to the published literature. The experimental study of Valero-Cuevas *et al*. [9] is the most complete data set that quantifies both thumb endpoint forces and muscle activations. However, pinch force was measured relative to the distal phalanx; normative lateral pinch protocols measure strength relative to a global frame [63, 77-79]. Similarly, the lateral pinch task defined by Valero-Cuevas *et al*. [9] was more restrained (participants had to limit off-axis forces to 17% of the normal force) than standard protocols (off-axis endpoint forces are not controlled). As noted in our previous publication [69], the choice of reference frame and how to replicate specific experimental conditions in simulations is not a trivial detail and can impact the interpretation of results (see discussion in [69]). We have previously successfully replicated the experiments (including the reference frame and all constraints) from [9], with the same thumb model implemented in a different computational environment [10, 69]. Here, we simulated pinch strength relative to the global frame.

Maximum lateral pinch force from our simulations (which incorporated the experimentally imposed restraint on pinch force direction) compared well to the magnitude of the lateral pinch force reported in [9], and the simulated forces were produced via muscle coordination patterns that generally fell within the variability of the EMG data from that study (Fig. 5). Given the smaller force magnitudes observed by [9] from normative pinch strength data reported for healthy adult males between ages of 20 and 24 (96.1-115.6N) [57, 80-82], our lateral pinch simulations were weaker than these normative data. Furthermore, our predicted activations were dissimilar to EMG data reported during normative pinch strength testing, which document >70% of full activation for APB, EPB, EPL and APL [80] (see Fig. 5 for our predicted activations). Simulated pinch strength only increased slightly (∼5N) and muscle coordination patterns did not change considerably when the restraint on pinch force direction was removed from our optimization. While the simulations presented here do not explain why the model is weaker than normative pinch strength, to our knowledge, no prior simulation study [1, 9, 10, 65, 69] has been able to replicate the pinch forces magnitudes reported in normative studies of adults. Additional experimental and simulation work will be required to understand why pinch simulations remain weaker than normative pinch strength. An important direction that may shed insight into this discrepancy is to better characterization of joint motion during pinch force production. For example, prior work has shown that thumb endpoint force is sensitive to joint posture [10, 81]; however, most experimental studies of pinch strength either don’t record thumb posture or set an initial posture without controlling or recording changes to posture during force production. Additionally, how the endpoint force is modeled in the simulations may influence pinch force production. Here, we adopted the current standard which is to model pinch force with a point force constraint, but future work could implement the elastic foundation contract forces used for grip strength to evaluate whether these choices influence results.

### C. Performance of model for active and passive hand motion

In general, timing and peaks of predicted activations from the active hand opening simulations agreed with the normalized EMG signals, indicating that our model can be used to predict reasonable activations for functional kinematic tasks. The largest discrepancy between the simulations and the EMG data (EDC at the start of the motion) appears to stem from the omission of the extensor mechanism from our model. Experimental and modeling studies have shown that the extensor mechanism plays an important role in transmission of force across the interphalangeal joints [82], coordination of finger movements [83, 84], and determination of both muscle and joint contact forces [24]. Because our model does not include an extensor mechanism, the intrinsics cannot contribute to PIP and DIP extension as they do *in vivo*, and EDC is the only extensor crossing the PIP and DIP joints. Thus, with our model, EDC may need to take on larger activations than measured in our participants to extend their fingers. Additionally, by omitting the extensor mechanism, 100% of the forces produced by the extrinsic finger extensors are transmitted across both interphalangeal joints. However, experimental work has shown that only ∼60% of the force transmitted through the central slip of the extensor mechanism was transmitted through the terminal slip to the distal interphalangeal joint [82]. Thus, without the extensor mechanism, it is likely that the resultant extensor torque at the interphalangeal joint is artifactually high, given that all the EDC force is transmitted across the PIP and DIP joint, which may explain why the reserve actuators for these joints needed to apply flexion torques to simulate hand opening. We anticipate that future work to better characterize and model the extensor mechanism will further improve predicted muscle activations during kinematic tasks.

For the passive hand opening simulation, the model displayed coupled motion between the wrist and digits mimicking tenodesis, indicating that our model displays appropriate motion in the absence of active forces. While the model displayed the passive motion associated with tenodesis, the individual fingers moved through the range of motion with different trajectories which is dissimilar to experimental kinematic of tenodesis with active wrist motion [26]. In addition to differences in muscle activation between the experimental and simulation study, assumptions in how passive joint moments were implemented may also contribute to the large variability in kinematics between the digits. Because the passive moments at the PIP and DIP joint of the middle, ring, and little finger have not been quantified, these passive properties were implemented through scaling passive moments measured at the index finger; the tendon slack lengths for the extrinsic muscles of these digits were computed from equation 1. On the other hand, passive joint moments for all the joints of the index finger have been previously quantified [25, 35], and prior work from our lab optimized the tendon slack lengths for the extrinsic finger muscles of the finger to better represent the passive joint moments measured experimentally [6]. This difference in the implementation of passive joint properties likely contributed to the notable differences in passive motion between the index finger and the other digits. Additionally, the skin between the fingers can create a resistance force between digits; this interconnected passive force has been modeled in other work as ligaments [13], but was not modeled here. Future work to better characterize and model both passive joint moments and skin resistance forces between the digits would likely improve passive simulations with our musculoskeletal model.

### D. Limitations of this work and future directions for the field

As has been presented throughout this work, we have identified several limitations (e.g., the lack of consistent data sets describing grip and pinch force production, omitting the extensor mechanism in the model, and incomplete description of passive joint properties) that need to be addressed in future work. Many of these limitations stem from limitations in the available data sets to build and validate the model. When developing the model, we were required to make several assumptions about the active and passive force generating capacity of muscles (particularly for the middle, ring, and little finger) due to lack of data describing moment arms (MCP abduction, PIP and DIP flexion), *in vivo* muscle volume (intrinsic muscles), and passive joint moments (interphalangeal joints of middle, ring, little and thumb). In general, to develop the model in the absence of these data, we had to scale data describing the index finger to the other digits. These missing data are not trivial to collect, and without a foundational hand and wrist model to incorporate these data, there has not been a sufficient need to collect these data.

When validating our model, the primary barrier was a lack of consistent data sets describing all aspects of force production (grip and pinch). For example, we had to compare our grip simulations to multiple experimental studies since currently no data set describes grip force, distribution of finger forces, EMG, and joint posture. Collecting such a multifaceted data set will be challenging, but future work can use the model and initial simulation work to design experimental studies to guide data collection. For example, our current simulations did not display the same sensitivity to wrist posture. Also, our results suggest that a better understanding of co-activations that occur during grasping may improve simulation performance. Here, we only required co-activation between the individual compartments of FDS, FDP, and EDC; however, involuntary co-activation also occurs between separate muscles and not just between compartments of a singular muscle during force production [56]. Thus, characterizing and incorporating involuntary co-activations during grasping is an important direction for experimental work that could potentially improve simulation performance and increase sensitivity to wrist posture.

In addition to the challenges from limited data sets needed to develop and validate the model, we have identified limitations in the technical implementation of biomechanical models that will need to be addressed in future work. Of highest priority, future work will need to move towards incorporating an extensor mechanism. The interconnected nature of the extensor mechanism makes it challenging to model in OpenSim. The intersecting bands of the extensor mechanism have been previously modeled as separate paths sharing via-points [8, 17]. However, this approach greatly increases the number of muscle-tendon actuators and requires multiple constraints to model force sharing amongst these paths, and thus was not used here for this initial implementation of our model. The current simulations highlight that future work to better characterize and implement the extensor mechanism is needed.

Lastly, in addition to those limitations, our musculoskeletal model was developed from multiple unique data sets. Prior studies highlight that unique data sets are not always mechanically consistent [10] and can lead to error in muscle force prediction [85]. For example, in the current work, developing the model from multiple data sets may have limited the sensitivity to wrist posture during grasping in the radial/ulnar direction since wrist strength of our model was set to match flexion/extension strength from a study that did not simultaneously measure wrist strength in radial/ulnar deviation [47]. The model has slightly greater radial/ulnar deviation capacity than reported in other studies in the literature [70]. Future work should compare simulation performance for these functional tasks with models developed from consistent data sets [13, 17].

## VII. Conclusion

We developed an open-source musculoskeletal model that includes the wrist, all digits and muscles of the hand, and passive joint properties for each flexion/extension DOF. This is the first open-source model of the hand and wrist to be implemented and evaluated during both functional kinetic and kinematic tasks. To our knowledge, this is the first implementation of an optimal control theory framework to predict both maximal grip strength and lateral pinch force using a muscle-driven biomechanical model. Overall, the model’s maximum grip force production was comparable to grip force and force distribution amongst the digits reported in the literature for healthy young adults. Lateral pinch strength simulated dynamically, using this optimal control theory framework was similar to previous simulations that use similar (or the same) thumb models under more constrained and static conditions. Simulated lateral pinch strength fell within variability of pinch strength data when off-axis forces are limited. This simulation framework provides the ability for future studies to evaluate how potential surgical and rehabilitation interventions influence clinical outcomes while requiring minimal experimental data as a simulation input. Additionally, we simulated active hand opening and tenodesis grasp and release to evaluate the model’s ability to simulate coordinated hand movements. While evaluations of these coordinated kinematic hand movements were less extensive than the kinetic functional tasks, our active and passive hand opening simulations predicted reasonable activations and demonstrated passive motion mimicking tenodesis, respectively. Overall, this open-source model and simulation tutorials provides a solid foundation for future work simulating coordinated kinematic functional tasks to build upon. Overall, our simulation results suggest that incorporating a model of the extensor mechanism, developing a better understand of muscle coordination during functional tasks, and better characterizing passive structures would further improve simulation outcomes.

## Acknowledgment

We would like to thank all past members of the ARMS Lab, especially James Buffi whose work defined the kinematic models of digits 3, 4, and 5 and established the calibration methods we used to link the model and the Cyberglove recordings. We would also like to thank Vikram Darbhe, Jorie Budzikowski, Maggie McDonough, Martin Seyres, and Morgan Dalman for their helpful input on the written tutorials (available on simtk.org) intended to enable others to replicate the simulations described here.

## References

[1] J. A. Nichols, M. S. Bednar, S. J. Wohlman, and W. M. Murray, “Connecting the wrist to the hand: A simulation study exploring changes in thumb-tip endpoint force following wrist surgery,” (in eng), J Biomech, vol. 58, pp. 97–104, Jun 14 2017, doi: 10.1016/j.jbiomech.2017.04.024.

[2] A. J. Barry, W. M. Murray, and D. G. Kamper, “Development of a dynamic index finger and thumb model to study impairment,” (in eng), J Biomech, vol. 77, pp. 206–210, Aug 22 2018, doi: 10.1016/j.jbiomech.2018.06.017.

[3] B. Goislard de Monsabert, L. Vigouroux, D. Bendahan, and E. Berton, “Quantification of finger joint loadings using musculoskeletal modelling clarifies mechanical risk factors of hand osteoarthritis,” (in eng), Med Eng Phys, vol. 36, no. 2, pp. 177–84, Feb 2014, doi: 10.1016/j.medengphy.2013.10.007.

[4] M. M. Adamczyk and P. E. Crago, “Simulated feedforward neural network coordination of hand grasp and wrist angle in a neuroprosthesis,” (in eng), IEEE Trans Rehabil Eng, vol. 8, no. 3, pp. 297–304, Sep 2000, doi: 10.1109/86.867871.

[5] B. I. Binder-Markey, J. P. A. Dewald, and W. M. Murray, “The Biomechanical Basis of the Claw Finger Deformity: A Computational Simulation Study,” (in eng), J Hand Surg Am, vol. 44, no. 9, pp. 751–761, Sep 2019, doi: 10.1016/j.jhsa.2019.05.007.

[6] B. I. Binder-Markey and W. M. Murray, “Incorporating the length-dependent passive-force generating muscle properties of the extrinsic finger muscles into a wrist and finger biomechanical musculoskeletal model,” (in eng), J Biomech, vol. 61, pp. 250–257, Aug 16 2017, doi: 10.1016/j.jbiomech.2017.06.026.

[7] A. Esteki and J. M. Mansour, “A dynamic model of the hand with application in functional neuromuscular stimulation,” (in eng), Ann Biomed Eng, vol. 25, no. 3, pp. 440–51, May-Jun 1997, doi: 10.1007/bf02684185.

[8] A. R. MacIntosh and P. J. Keir, “An open-source model and solution method to predict co-contraction in the finger,” (in eng), Comput Methods Biomech Biomed Engin, vol. 20, no. 13, pp. 1373–1381, Oct 2017, doi: 10.1080/10255842.2017.1364732.

[9] F. J. Valero-Cuevas, M. E. Johanson, and J. D. Towles, “Towards a realistic biomechanical model of the thumb: the choice of kinematic description may be more critical than the solution method or the variability/uncertainty of musculoskeletal parameters,” (in eng), J Biomech, vol. 36, no. 7, pp. 1019–30, Jul 2003, doi: 10.1016/s0021-9290(03)00061-7.

[10] S. J. Wohlman and W. M. Murray, “Bridging the gap between cadaveric and in vivo experiments: a biomechanical model evaluating thumb-tip endpoint forces,” (in eng), J Biomech, vol. 46, no. 5, pp. 1014–20, Mar 15 2013, doi: 10.1016/j.jbiomech.2012.10.044.

[11] K. A. Hao and J. A. Nichols, “Simulating finger-tip force using two common contact models: Hunt-Crossley and elastic foundation,” (in eng), J Biomech, vol. 119, p. 110334, Apr 15 2021, doi: 10.1016/j.jbiomech.2021.110334.

[12] T. Ordonez Diaz and J. A. Nichols, “Anthropometric scaling of musculoskeletal models of the hand captures age-dependent differences in lateral pinch force,” (in eng), J Biomech, vol. 123, p. 110498, Jun 23 2021, doi: 10.1016/j.jbiomech.2021.110498.

[13] L. Engelhardt et al., “A new musculoskeletal AnyBody(tm) detailed hand model,” (in eng), Comput Methods Biomech Biomed Engin, pp. 1–11, Dec 10 2020, doi: 10.1080/10255842.2020.1851367.

[14] B. Goislard de Monsabert, J. Rossi, E. Berton, and L. Vigouroux, “Quantification of hand and forearm muscle forces during a maximal power grip task,” (in eng), Med Sci Sports Exerc, vol. 44, no. 10, pp. 1906–16, Oct 2012, doi: 10.1249/MSS.0b013e31825d9612.

[15] J. H. Lee, D. S. Asakawa, J. T. Dennerlein, and D. L. Jindrich, “Finger muscle attachments for an OpenSim upper-extremity model,” (in eng), PLoS One, vol. 10, no. 4, p. e0121712,2015, doi: 10.1371/journal.pone.0121712.

[16] J. Ma’touq, T. Hu, and S. Haddadin, “A validated combined musculotendon path and muscle-joint kinematics model for the human hand,” (in eng), Comput Methods Biomech Biomed Engin, vol. 22, no. 7, pp. 727–739, May 2019, doi: 10.1080/10255842.2019.1588256.

[17] M. Mirakhorlo, N. Van Beek, M. Wesseling, H. Maas, H. E. J. Veeger, and I. Jonkers, “A musculoskeletal model of the hand and wrist: model definition and evaluation,” (in eng), Comput Methods Biomech Biomed Engin, pp. 1–10, Sep 26 2018, doi: 10.1080/10255842.2018.1490952.

[18] J. Rossi, B. Goislard De Monsabert, E. Berton, and L. Vigouroux, “Handle Shape Affects the Grip Force Distribution and the Muscle Loadings During Power Grip Tasks,” (in eng), J Appl Biomech, vol. 31, no. 6, pp. 430–8, Dec 2015, doi: 10.1123/jab.2014-0171.

[19] F. E. Zajac, “Muscle coordination of movement: a perspective,” (in eng), J Biomech, vol. 26 Suppl 1, pp. 109–24, 1993, doi: 10.1016/0021-9290(93)90083-q.

[20] S. A. Kautz, R. R. Neptune, and F. E. Zajac, “General coordination principles elucidated by forward dynamics: minimum fatique does not explain muscle excitation in dynamic tasks,” (in eng), Motor Control, vol. 4, no. 1, pp. 75–80; discussion 97-116, Jan 2000, doi: 10.1123/mcj.4.1.75.

[21] M. G. Pandy, F. E. Zajac, E. Sim, and W. S. Levine, “An optimal control model for maximum-height human jumping,” (in eng), J Biomech, vol. 23, no. 12, pp. 1185–98, 1990, doi: 10.1016/0021-9290(90)90376-e.

[22] Z. M. Li, L. Kuxhaus, J. A. Fisk, and T. H. Christophel, “Coupling between wrist flexion-extension and radial-ulnar deviation,” (in eng), Clin Biomech (Bristol, Avon), vol. 20, no. 2, pp. 177–83, Feb 2005, doi: 10.1016/j.clinbiomech.2004.10.002.

[23] H. Moritomo, E. P. Apergis, G. Herzberg, F. W. Werner, S. W. Wolfe, and M. Garcia-Elias, “2007 IFSSH committee report of wrist biomechanics committee: biomechanics of the so-called dart-throwing motion of the wrist,” (in eng), J Hand Surg Am, vol. 32, no. 9, pp. 1447–53, Nov 2007, doi: 10.1016/j.jhsa.2007.08.014.

[24] D. Hu, D. Howard, and L. Ren, “Biomechanical analysis of the human finger extensor mechanism during isometric pressing,” (in eng), PLoS One, vol. 9, no. 4, p. e94533. 2014, doi: 10.1371/journal.pone.0094533.

[25] D. G. Kamper, T. George Hornby, and W. Z. Rymer, “Extrinsic flexor muscles generate concurrent flexion of all three finger joints,” (in eng), J Biomech, vol. 35, no. 12, pp. 1581–9, Dec 2002, doi: 10.1016/s0021-9290(02)00229-4.

[26] F. C. Su, Y. L. Chou, C. S. Yang, G. T. Lin, and K. N. An, “Movement of finger joints induced by synergistic wrist motion,” (in eng), Clin Biomech (Bristol, Avon), vol. 20, no. 5, pp. 491–7, Jun 2005, doi: 10.1016/j.clinbiomech.2005.01.002.

[27] S. L. Delp et al., “OpenSim: open-source software to create and analyze dynamic simulations of movement,” (in eng), IEEE Trans Biomed Eng, vol. 54, no. 11, pp. 1940–50, Nov 2007, doi: 10.1109/tbme.2007.901024.

[28] D. Blana, E. K. Chadwick, A. J. van den Bogert, and W. M. Murray, “Real-time simulation of hand motion for prosthesis control,” (in eng), Comput Methods Biomech Biomed Engin, vol. 20, no. 5, pp. 540–549, Apr 2017, doi: 10.1080/10255842.2016.1255943.

[29] J. H. Buffi, J. J. Crisco, and W. M. Murray, “A method for defining carpometacarpal joint kinematics from three-dimensional rotations of the metacarpal bones captured in vivo using computed tomography,” (in eng), J Biomech, vol. 46, no. 12, pp. 2104–8, Aug 9 2013, doi: 10.1016/j.jbiomech.2013.05.019.

[30] J. H. Buffi, J. L. Sancho Bru, J. J. Crisco, and W. M. Murray, “Evaluation of hand motion capture protocol using static computed tomography images: application to an instrumented glove,” (in eng), J Biomech Eng, vol. 136, no. 12, p. 124501, Dec 2014, doi: 10.1115/1.4028521.

[31] K. R. Holzbaur, W. M. Murray, and S. L. Delp, “A model of the upper extremity for simulating musculoskeletal surgery and analyzing neuromuscular control,” (in eng), Ann Biomed Eng, vol. 33, no. 6, pp. 829–40, Jun 2005, doi: 10.1007/s10439-005-3320-7.

[32] K. R. Saul et al., “Benchmarking of dynamic simulation predictions in two software platforms using an upper limb musculoskeletal model,” (in eng), Comput Methods Biomech Biomed Engin, vol. 18, no. 13, pp. 1445–58, 2015, doi: 10.1080/10255842.2014.916698.

[33] R. Degeorges, S. Laporte, E. Pessis, D. Mitton, J. N. Goubier, and F. Lavaste, “Rotations of three-joint fingers: a radiological study,” (in eng), Surg Radiol Anat, vol. 26, no. 5, pp. 392–8, Oct 2004, doi: 10.1007/s00276-004-0244-0.

[34] M. Domalain, L. Vigouroux, and E. Berton, “Determination of passive moment-angle relationships at the trapeziometacarpal joint,” (in eng), J Biomech Eng, vol. 132, no. 7, p. 071009, Jul 2010, doi: 10.1115/1.4001397.

[35] J. S. Knutson, K. L. Kilgore, J. M. Mansour, and P. E. Crago, “Intrinsic and extrinsic contributions to the passive moment at the metacarpophalangeal joint,” (in eng), J Biomech, vol. 33, no. 12, pp. 1675–81, Dec 2000, doi: 10.1016/s0021-9290(00)00159-7.

[36] Z. M. Li, G. Davis, N. P. Gustafson, and R. J. Goitz, “A robot-assisted study of intrinsic muscle regulation on proximal interphalangeal joint stiffness by varying metacarpophalangeal joint position,” (in eng), J Orthop Res, vol. 24, no. 3, pp. 407–15, Mar 2006, doi: 10.1002/jor.20046.

[37] B. I. Binder-Markey, W. M. Murray, and J. Dewald, “Passive properties of the wrist and fingers following chronic hemiparetic stroke: interlimb comparisons in persons with and without a clinical treatment history that includes Botulinum Neurotoxin,” Frontiers in Neurology, vol. 12, p. 1290, 2021.

[38] “OpenSim::CoordinateLimitForce Class Reference,” ed.

[39] S. Koh, W. L. Buford, Jr., C. R. Andersen, and S. F. Viegas, “Intrinsic muscle contribution to the metacarpophalangeal joint flexion moment of the middle, ring, and small fingers,” (in eng), J Hand Surg Am, vol. 31, no. 7, pp. 1111–7, Sep 2006, doi: 10.1016/j.jhsa.2006.03.003.

[40] K. N. An, Y. Ueba, E. Y. Chao, W. P. Cooney, and R. L. Linscheid, “Tendon excursion and moment arm of index finger muscles,” (in eng), J Biomech, vol. 16, no. 6, pp. 419–25, 1983, doi: 10.1016/0021-9290(83)90074-x.

[41] W. L. Buford, Jr., S. Koh, C. R. Andersen, and S. F. Viegas, “Analysis of intrinsic-extrinsic muscle function through interactive 3-dimensional kinematic simulation and cadaver studies,” (in eng), J Hand Surg Am, vol. 30, no. 6, pp. 1267–75, Nov 2005, doi: 10.1016/j.jhsa.2005.06.019.

[42] A. Synek and D. H. Pahr, “The effect of the extensor mechanism on maximum isometric fingertip forces: A numerical study on the index finger,” (in eng), J Biomech, vol. 49, no. 14, pp. 3423–3429, Oct 3 2016, doi: 10.1016/j.jbiomech.2016.09.004.

[43] M. Millard, T. Uchida, A. Seth, and S. L. Delp, “Flexing computational muscle: modeling and simulation of musculotendon dynamics,” (in eng), J Biomech Eng, vol. 135, no. 2, p. 021005, Feb 2013, doi: 10.1115/1.4023390.

[44] M. D. Jacobson, R. Raab, B. M. Fazeli, R. A. Abrams, M. J. Botte, and R. L. Lieber, “Architectural design of the human intrinsic hand muscles,” (in eng), J Hand Surg Am, vol. 17, no. 5, pp. 804–9, Sep 1992, doi: 10.1016/0363-5023(92)90446-v.

[45] F. D. Kerkhof, T. van Leeuwen, and E. E. Vereecke, “The digital human forearm and hand,” (in eng), J Anat, vol. 233, no. 5, pp. 557–566, Nov 2018, doi: 10.1111/joa.12877.

[46] K. R. Holzbaur, W. M. Murray, G. E. Gold, and S. L. Delp, “Upper limb muscle volumes in adult subjects,” (in eng), J Biomech, vol. 40, no. 4, pp. 742–9, 2007, doi: 10.1016/j.jbiomech.2006.11.011.

[47] K. R. Holzbaur, S. L. Delp, G. E. Gold, and W. M. Murray, “Moment-generating capacity of upper limb muscles in healthy adults,” (in eng), J Biomech, vol. 40, no. 11, pp. 2442–9, 2007, doi: 10.1016/j.jbiomech.2006.11.013.

[48] S. J. Wohlman, “Understanding the dynamics of thumb-tip force generation through integration of simulation and experimental methods,” Northwestern University, Evanston IL (Doctor of Philosophy), 2015.

[49] L. Blankevoort, J. H. Kuiper, R. Huiskes, and H. J. Grootenboer, “Articular contact in a three-dimensional model of the knee,” (in eng), J Biomech, vol. 24, no. 11, pp. 1019–31, 1991, doi: 10.1016/0021-9290(91)90019-j.

[50] M. A. Sherman, A. Seth, and S. L. Delp, “Simbody: multibody dynamics for biomedical research,” (in eng), Procedia IUTAM, vol. 2, pp. 241–261, 2011, doi: 10.1016/j.piutam.2011.04.023.

[51] E. Fess, “Clinical assessment recommendations,” American society of hand therapists, pp. 6–8, 1981.

[52] J. C. Firrell and G. M. Crain, “Which setting of the dynamometer provides maximal grip strength?,” (in eng), J Hand Surg Am, vol. 21, no. 3, pp. 397–401, May 1996, doi: 10.1016/s0363-5023(96)80351-0.

[53] S. E. Tomlinson, R. Lewis, and M. J. Carré, “The effect of normal force and roughness on friction in human finger contact,” Wear, vol. 267, no. 5, pp. 1311–1318, 2009/06/15/ 2009, doi: https://doi.org/10.1016/j.wear.2008.12.084.

[54] C. Li, G. Guan, R. Reif, Z. Huang, and R. K. Wang, “Determining elastic properties of skin by measuring surface waves from an impulse mechanical stimulus using phase-sensitive optical coherence tomography,” (in eng), J R Soc Interface, vol. 9, no. 70, pp. 831–41, May 7 2012, doi: 10.1098/rsif.2011.0583.

[55] M. Pawlaczyk, M. Lelonkiewicz, and M. Wieczorowski, “Age-dependent biomechanical properties of the skin,” (in eng), Postepy Dermatol Alergol, vol. 30, no. 5, pp. 302–6, Oct 2013, doi: 10.5114/pdia.2013.38359.

[56] J. A. Birdwell, L. J. Hargrove, T. A. Kuiken, and R. F. Weir, “Activation of individual extrinsic thumb muscles and compartments of extrinsic finger muscles,” (in eng), J Neurophysiol, vol. 110, no. 6, pp. 1385–92, Sep 2013, doi: 10.1152/jn.00748.2012.

[57] S. W. O’Driscoll, E. Horii, R. Ness, T. D. Cahalan, R. R. Richards, and K. N. An, “The relationship between wrist position, grasp size, and grip strength,” (in eng), J Hand Surg Am, vol. 17, no. 1, pp. 169–77, Jan 1992, doi: 10.1016/0363-5023(92)90136-d.

[58] Y. K. Kong, K. S. Lee, D. M. Kim, and M. C. Jung, “Individual finger contribution in submaximal voluntary contraction of gripping,” (in eng), Ergonomics, vol. 54, no. 11, pp. 1072–80, Nov 2011, doi: 10.1080/00140139.2011.620176.

[59] J. P. Mogk and P. J. Keir, “The effects of posture on forearm muscle loading during gripping,” (in eng), Ergonomics, vol. 46, no. 9, pp. 956–75, Jul 15 2003, doi: 10.1080/0014013031000107595.

[60] J. A. Balogun, S. A. Adenlola, and A. A. Akinloye, “Grip strength normative data for the harpenden dynamometer,” (in eng), J Orthop Sports Phys Ther, vol. 14, no. 4, pp. 155–60, 1991, doi: 10.2519/jospt.1991.14.4.155.

[61] J. Y. Hogrel, “Grip strength measured by high precision dynamometry in healthy subjects from 5 to 80 years,” (in eng), BMC Musculoskelet Disord, vol. 16, p. 139, Jun 10 2015, doi: 10.1186/s12891-015-0612-4.

[62] S. L. Wong, “Grip strength reference values for Canadians aged 6 to 79: Canadian Health Measures Survey, 2007 to 2013,” (in eng), Health Rep, vol. 27, no. 10, pp. 3–10, Oct 19 2016.

[63] V. Mathiowetz, N. Kashman, G. Volland, K. Weber, M. Dowe, and S. Rogers, “Grip and pinch strength: normative data for adults,” (in eng), Arch Phys Med Rehabil, vol. 66, no. 2, pp. 69–74, Feb 1985.

[64] M. E. Johanson, M. A. James, and S. R. Skinner, “Forearm muscle activation during power grip and release,” (in eng), J Hand Surg Am, vol. 23, no. 5, pp. 938–44, Sep 1998, doi: 10.1016/s0363-5023(98)80177-9.

[65] D. C. McFarland, J. A. Nichols, M. S. Bednar, S. J. Wohlman, and W. M. Murray, “Corrigendum to “Connecting the wrist to the hand: A simulation study exploring changes in thumb-tip endpoint force following wrist surgery” [J. Biomech. 58 (2017) 97–104],” (in eng), J Biomech, p. 110859, Nov 20 2021, doi: 10.1016/j.jbiomech.2021.110859.

[66] “How Static Optimization Works.” https://simtk-confluence.stanford.edu/display/OpenSim/How+Static+Optimization+Works (accessed 8/11/2021.

[67] M. Melzner, L. Engelhardt, U. Simon, and S. Dendorfer, “Electromyography-Based Validation of a Musculoskeletal Hand Model,” (in eng), J Biomech Eng, vol. 144, no. 2, Feb 1 2022, doi: 10.1115/1.4052115.

[68] D. G. Thelen, F. C. Anderson, and S. L. Delp, “Generating dynamic simulations of movement using computed muscle control,” (in eng), J Biomech, vol. 36, no. 3, pp. 321–8, Mar 2003, doi: 10.1016/s0021-9290(02)00432-3.

[69] D. C. McFarland, S. J. Wohlman, and W. M. Murray, “Corrigendum to “Bridging the gap between cadaveric and in vivo experiments: A biomechanical model evaluating thumb-tip endpoint forces” [J. Biomech. 46(5) (2013) 1014–1020],” (in eng), J Biomech, p. 110858, Nov 19 2021, doi: 10.1016/j.jbiomech.2021.110858.

[70] S. L. Delp, A. E. Grierson, and T. S. Buchanan, “Maximum isometric moments generated by the wrist muscles in flexion-extension and radial-ulnar deviation,” (in eng), J Biomech, vol. 29, no. 10, pp. 1371–5, Oct 1996, doi: 10.1016/0021-9290(96)00029-2.

[71] M. Caumes, B. Goislard de Monsabert, H. Hauraix, E. Berton, and L. Vigouroux, “Complex couplings between joints, muscles and performance: the role of the wrist in grasping,” (in eng), Sci Rep, vol. 9, no. 1, p. 19357, Dec 18 2019, doi: 10.1038/s41598-019-55443-w.

[72] P. W. Fong and G. Y. Ng, “Effect of wrist positioning on the repeatability and strength of power grip,” (in eng), Am J Occup Ther, vol. 55, no. 2, pp. 212–6, Mar-Apr 2001, doi: 10.5014/ajot.55.2.212.

[73] F. T. Hazelton, G. L. Smidt, A. E. Flatt, and R. I. Stephens, “The influence of wrist position on the force produced by the finger flexors,” (in eng), J Biomech, vol. 8, no. 5, pp. 301–6, Sep 1975, doi: 10.1016/0021-9290(75)90082-2.

[74] J. C. Pryce, “The wrist position between neutral and ulnar deviation that facilitates the maximum power grip strength,” (in eng), J Biomech, vol. 13, no. 6, pp. 505–11, 1980, doi: 10.1016/0021-9290(80)90343-7.

[75] D. C. Ackland, Y. C. Lin, and M. G. Pandy, “Sensitivity of model predictions of muscle function to changes in moment arms and muscle-tendon properties: a Monte-Carlo analysis,” (in eng), J Biomech, vol. 45, no. 8, pp. 1463–71, May 11 2012, doi: 10.1016/j.jbiomech.2012.02.023.

[76] E. M. Arnold, S. R. Ward, R. L. Lieber, and S. L. Delp, “A model of the lower limb for analysis of human movement,” (in eng), Ann Biomed Eng, vol. 38, no. 2, pp. 269–79, Feb 2010, doi: 10.1007/s10439-009-9852-5.

[77] E. Fain and C. Weatherford, “Comparative study of millennials’ (age 20-34 years) grip and lateral pinch with the norms,” (in eng), J Hand Ther, vol. 29, no. 4, pp. 483–488, Oct-Dec 2016, doi: 10.1016/j.jht.2015.12.006.

[78] M. Mohammadian, A. Choobineh, A. Haghdoost, and N. Hasheminejad, “Normative data of grip and pinch strengths in healthy adults of Iranian population,” (in eng), Iran J Public Health, vol. 43, no. 8, pp. 1113–22, Aug 2014.

[79] S. Werle, J. Goldhahn, S. Drerup, B. R. Simmen, H. Sprott, and D. B. Herren, “Age-and gender-specific normative data of grip and pinch strength in a healthy adult Swiss population,” (in eng), J Hand Surg Eur Vol, vol. 34, no. 1, pp. 76–84, Feb 2009, doi: 10.1177/1753193408096763.

[80] F. D. Kerkhof, G. Deleu, P. D’Agostino, and E. E. Vereecke, “Subject-specific thumb muscle activity during functional tasks of daily life,” (in eng), J Electromyogr Kinesiol, vol. 30, pp. 131–6, Oct 2016, doi: 10.1016/j.jelekin.2016.06.009.

[81] C. M. Goehler and W. M. Murray, “The sensitivity of endpoint forces produced by the extrinsic muscles of the thumb to posture,” (in eng), J Biomech, vol. 43, no. 8, pp. 1553–9, May 28 2010, doi: 10.1016/j.jbiomech.2010.01.032.

[82] S. W. Lee, H. Chen, J. D. Towles, and D. G. Kamper, “Effect of finger posture on the tendon force distribution within the finger extensor mechanism,” (in eng), J Biomech Eng, vol. 130, no. 5, p. 051014, Oct 2008, doi: 10.1115/1.2978983.

[83] J. M. Landsmeer, “The anatomy of the dorsal aponeurosis of the human finger and its functional significance,” (in eng), Anat Rec, vol. 104, no. 1, pp. 31–44, May 1949, doi: 10.1002/ar.1091040105.

[84] J. N. Leijnse and C. W. Spoor, “Reverse engineering finger extensor apparatus morphology from measured coupled interphalangeal joint angle trajectories - a generic 2D kinematic model,” (in eng), J Biomech, vol. 45, no. 3, pp. 569–78, Feb 2 2012, doi: 10.1016/j.jbiomech.2011.11.002.

[85] B. Goislard De Monsabert, D. Edwards, D. Shah, and A. Kedgley, “Importance of Consistent Datasets in Musculoskeletal Modelling: A Study of the Hand and Wrist,” (in eng), Ann Biomed Eng, vol. 46, no. 1, pp. 71–85, Jan 2018, doi: 10.1007/s10439-017-1936-z.

